# In vivo, online label-free monitoring of heterogenous oxygen utilization during phototherapy with real-time ultrasound-guided photoacoustic imaging

**DOI:** 10.1101/2024.11.27.625759

**Authors:** Andrew Langley, Allison Sweeney, Ronak T. Shethia, Brooke Bednarke, Faizah Wulandana, Marvin Xavierselvan, Srivalleesha Mallidi

## Abstract

Understanding the tumor microenvironment, particularly the vascular density and the availability of oxygen, is key in individualizing treatment approaches and determining their efficacy. While there are many therapies including radiotherapy that are ineffective in hypoxic tumor microenvironments, here we demonstrate the heterogeneous oxygen consumption during photodynamic therapy (PDT), a non-invasive treatment method using localized light to activate a photosensitive drug in the presence of oxygen that has shown high effectiveness in the treatment of various types of tumors, including those presented in head and neck cancer (HNC) patients. While our previous work has demonstrated that blood oxygen saturation (StO_2_) mapped before and after treatment with ultrasound-guided photoacoustic imaging (US-PAI) can be used as a surrogate marker for the regionalized long-term efficacy of PDT, real-time monitoring of StO_2_ during PDT could provide additional insights on oxygen consumption and inform dose design for “on the spot” treatment decisions. Specifically, in this work, we integrated the US-PAI transducer probe with PDT light delivery fibers. We tested the setup on murine tumor models intravenously injected with liposomal benzoporphyrin derivative (BPD) photosensitizer at 0.5 mg/kg dose and photodynamic illumination at 100 and 400 mW/cm^2^ fluence rate. As expected, we observed with our US-PAI StO_2_ images that the rate of oxygen utilization increases when using a high fluence rate (HFR) light dose. Particularly in the higher fluence rate group, we observed StO_2_ reaching a minimum mid-light dose, followed by some degree of re-oxygenation. US-PAI added the advantage of spatial information to StO_2_ monitoring, which allowed us to match regions of re-oxygenation during therapy to retained vascular function with immunohistochemistry. Overall, our results have demonstrated the potential of US-PAI for applications in online dosimetry for cancer therapies such as PDT, using oxygen changes to detect regionalized physiological vascular response in the tumor microenvironment.

## Introduction

Hemodynamic measurements, including vascular density, perfusion, and blood oxygen saturation (StO_2_), have emerged as vital prognostic markers for evaluating the efficacy of various cancer treatments. ^1-3^ These measurements are crucial, due to the vital role played by the tumor vasculature in growth and proliferation. Amongst different types of cancer, head and neck cancer (HNC) presents unique challenges in treatment due to the proximity to functional tissues abundant with vasculature hence making the non-invasive study of tumor vascular dynamics particularly significant. Outcomes in HNC, which encompasses malignancies in the oral cavity, pharynx, or larynx, have not improved at the same rate as other cancer types despite widespread technological advancements.^4^ HNC is often linked to risk factors such as tobacco use, excessive alcohol consumption, and human papillomavirus infection, with a higher prevalence in regions with limited healthcare infrastructure.^5^ Treatment modalities for HNC, including surgical resection, chemotherapy, and radiation therapy, while each having distinct advantages and disadvantages, have not significantly improved the overall quality of life for patients. Surgical resection is effective for eliminating the lesion, but lack of cellular specificity necessitates precautionary tumor margin removal and needless loss of healthy tissue. Loss of healthy tissue in sensitive areas can diminish the patient’s quality of life by impeding jaw, neck, or tongue function critical to speech and swallowing, as well as self-image from cosmetic damage.^6^ Chemotherapy provides a systemic treatment approach but typically involves drugs with off-target effects leading to undesirable patient side effects^7^ while radiation therapy exposes the patient to harmful ionizing radiation with relatively poor target specificity, damaging surrounding healthy tissues.^8^

In recent years, photodynamic therapy (PDT) has gained prominence as a specialized, cost-effective treatment for solid localized lesions in HNC,^9,10^ offering the benefits of minimal scarring, high selectivity, and low systemic toxicity through the use of light-activated drugs.^9-13^ PDT is highly selective as it requires spatially localized optical irradiation. In addition, preferential drug accumulation can dually minimize inadvertent treatment of healthy tissue.^14^ During PDT, a drug known as a photosensitizer (PS) is administered to the patient and the tumor is illuminated at a PS-specific wavelength. The excited PS undergoes intersystem crossing and reacts with oxygen to generate cytotoxic reactive oxygen species (ROS) that can destroy tissue through tumor cell death or vascular destruction, depending on the PS, mode of delivery, and drug-light interval.^13-16^ A major limitation of PDT is its notoriously complex dosimetry arising from the dynamic interplay between PS distribution, endogenous oxygen content, light dose, and tissue optical properties that will vary from patient to patient.^16-20^ While the physician has control over the administration of light and PS, the endogenous oxygen and micro-localization of PS available for photodynamic action are dependent on vascular density, metabolic activity, and tissue perfusion, all of which can be heterogeneous in tumor tissue.^16,21,22^ To account for the multitude of known factors that play into PDT dosimetry, there is a need for online monitoring of PDT to inform personalized dosimetry, which is elegantly detailed by Wilson et al.^16^ Effective PDT with vascular targeting PS has been shown to deplete tissue oxygen content over time, as tumor cells are starved of nutrients.^23^ Mallidi et al. found that tumor oxygen saturation (StO_2_) changes were indicative of PDT efficacy starting 24 hours post-PDT with liposomal benzoporphyrin derivative (BPD) PS.^23^ Wang et al found that StO_2_ measurements taken immediately after PDT with the Photofrin PS are sufficient to predict long-term efficacy.^24^ With different PSs having unique mechanism of action, it is unsurprising that the critical time point to predict PDT outcome also varies. The capability to identify non-responsive tumors within the hours after PDT is invaluable to shift the therapeutic approach early. Moreover, real-time monitoring may allow clinicians to classify sub-optimal therapeutic conditions during treatment and modulate the PDT for a more favorable, patient-tailored outcome. Light dose fluence rate and fractionation period are factors that have been shown to increase the effectiveness of PDT and can be easily tuned in real-time in response to monitoring.^16,19,25,26^ Given the oxygen-dependent nature of PDT’s mechanism, monitoring changes vascular hemodynamics (example density and oxygen saturation) can better aid clinicians in tailoring PDT parameters to enhance its therapeutic efficacy while minimizing damage to surrounding healthy tissues.

Several preclinical and clinical imaging techniques have been utilized to measure hemodynamics during PDT. Real-time blood flow measurements have been demonstrated during PDT with laser Doppler imaging^27^ and diffuse correlation spectroscopy (DCS),^28,29^ both of which provide insight into the physiological vascular response to PDT. However, both techniques are limited to a point-source measurement while laser Doppler imaging has the added disadvantages of the probe being invasive and easily saturated by PDT irradiation.^27,28^ Power Doppler ultrasound can provide non-invasive spatially resolved blood flow measurements but is currently hindered by lack of information on oxygenation,^28,30^ as are DCS and laser Doppler imaging. Blood flow provides inferential information on StO_2_ from the influx on oxygenated blood, but lacks the influence of varying oxygen utilization from ROS formation. Real-time oxygen monitoring during PDT has been successfully demonstrated with electrode measurements of pO_2_,^19,27^ and provides a direct measurement of molecular oxygen, but the probe is invasive, easily saturated by PDT irradiation, and the measurement is limited to a point source. Previous work has shown the utility of StO_2_ as a surrogate marker for tumor oxygen.^23,31^ StO_2_ has been previously monitored in real-time during PDT with diffuse reflectance spectroscopy, but once again this measurement lacks spatial information.^32^ Non-spatially-resolved hemodynamic measurements integrate values over a region and fail to capture the heterogeneity of the tumor vascular microenvironment. Blood oxygenation level-dependent contrast functional magnetic resonance imaging (BOLD fMRI), which provides spatial resolution, has also been used for real-time PDT monitoring of StO_2_ but is only sensitive to the paramagnetic response of deoxyhemoglobin (HbD), not oxyhemoglobin (HbO).^33^ BOLD fMRI would correlate with StO_2_ with constant hemoglobin amounts, but this is not the case during PDT due to established blood flow variation. BOLD fMRI signal must therefore be corrected by a secondary method to measure total hemoglobin. Additionally, BOLD fMRI is bulky, expensive, requires specialized training, and is overall impractical for PDT monitoring that is often conducted in infrastructure-limited countries as an inexpensive and portable cancer therapy to tackle health disparities, particularly for HNC.^5,9^

There has been recent interest in applications of photoacoustic (PA) imaging of StO_2_ to PDT. PA imaging is a 3D, non-ionizing imaging technique with high spatial resolution and contrast based on optical absorption.^34,35^ When a chromophore is irradiated with nanosecond pulsed light, it undergoes rapid thermoelastic expansion and contraction and creates pressure waves detectable by an ultrasound (US) transducer. Multiwavelength PA imaging can generate StO_2_ images of tissue based on the absorption differences between HbO and HbD, and functional hemodynamic information from PA imaging is easily co-registered with structural information from B-mode US due to their common hardware.^23,36-38^ Additionally, US combined with PA imaging (termed as US-PAI) can be incorporated into inexpensive and portable systems that are easy to operate, particularly with light-emitting diode illumination.^37,39^ For these reasons, US-PAI has been shown by our group and others to be a valuable research tool for monitoring the efficacy of PDT longitudinally,^23,40-42^ but until now, real-time PDT monitoring with US-PAI has been a mechanical challenge. We have developed a 3D-printed custom four-way PDT optical fiber holding attachment that easily attaches to a linear-array transducer (LAT) and angles the PDT fibers at the narrow transducer-to-skin gap for even tumor illumination from four sides. This design has allowed us to monitor PDT with US-PAI in real-time, *in vivo*, and we have demonstrated that the spatial resolving capabilities of US-PAI can be applied to real-time online monitoring of StO_2_ as a surrogate marker for oxygen utilization and vascular response during PDT.

## Materials and Methods

### Murine tumor model

All animal procedures were performed in accordance with the Institutional Animal Care and Use Committee (IACUC) at Tufts University. FaDu cells (human hypopharyngeal squamous cell carcinoma HNC, ATCC) were cultured in Dulbecco’s Modified Eagle’s Medium supplemented with 10% fetal bovine serum (FBS, Gibco) and 1% Penicillin and Streptomycin, (Corning) at 37°C and 5% CO_2_. Cells were passaged and used in experiments while in the exponential growth phase. Cells were trypsinized and suspended in a 1:1 mixture of phosphate-buffered saline (PBS) and Matrigel (Corning Inc.). Male homozygous Foxn1^nu^ mice (The Jackson Laboratory, Bar Harbor, ME) were sedated with 2% isoflurane and 5 million cells in 100 µL of the cell suspension were implanted subcutaneously into the lower back. The timeline for experiments is depicted in Fig. 1a. Following implantation, mouse body weight and tumor volume were monitored every other day. Tumors were allowed to grow for 8-10 days and progression was monitored with calipers utilizing the ellipsoid equation until reaching a volume of 100 mm^3^ (diameter ∼7-9 mm).^43^

**Figure 1.**
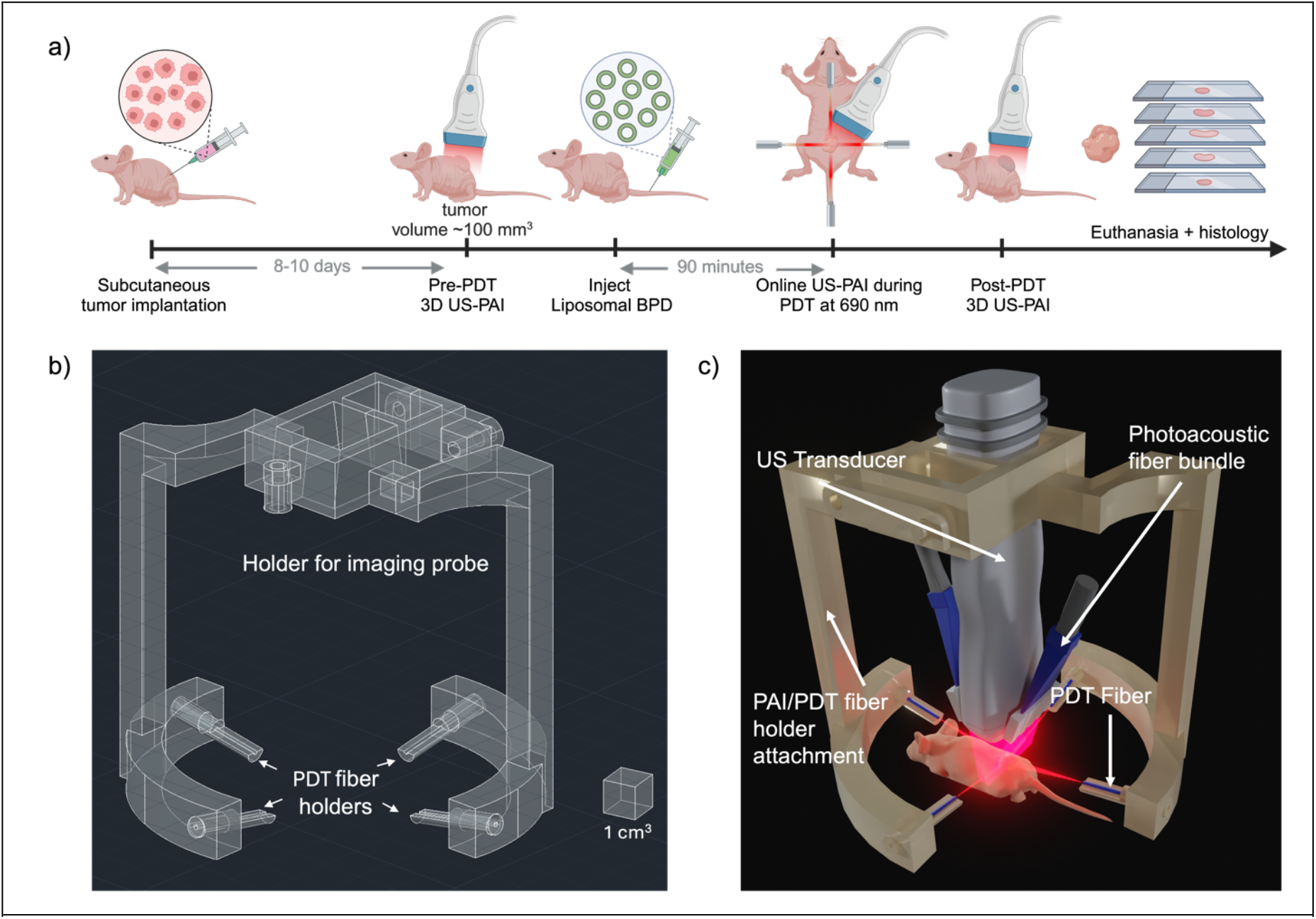
a) Experiment timeline for US-PAI, PDT treatment monitoring, and euthanasia of murine tumor models. Image was created using Biorender b) Computer aided design of the PDT fiber holder transducer attachment. c) Render of the fiber holder (3D printed with polylactic acid filament) attached to the transducer and PDT fibers.

Mice were anesthetized with isoflurane (2-2.5% for induction and 1.5-2% for maintenance) via nose cone on a heating pad for all procedures. Immediately following PDT and imaging procedures (described in detail below), pimonidazole (Hypoxyprobe, Inc.) was administered at a dose of 60 mg/kg via tail vein injection 60 minutes prior to euthanasia, and tomato lectin (TL) (Vector Laboratories) was administered at a dose of 50 mg 10 minutes prior to sacrifice. Euthanasia was performed via cervical dislocation and the tumors were extracted and frozen in Optimal Cutting Temperature compound (Tissue-Tek) at -80°C.

### Optical Fiber Mount Fabrication

The PDT fiber mount was customized for attachment to the Vevo LAZR-X MX250S transducer and necessitates four PDT optical fibers to evenly illuminate the tumor from the cranial, caudal, and both lateral sides of the mouse. The PDT fiber mount was designed in AutoCAD (AutoDesk), shown in Fig. 1b and 3D printed with clear polylactic acid filament. The design consisted of a latched clamp with two descending arms attached to a segmented elliptical ring (62 × 57 mm) fixed with four PDT optical fiber holders. All four fiber holders were angled at 10º from the elliptical plane, which allowed for more uniform illumination of the entire tumor. The length of the descending arms and geometry of the fiber holders allowed the PDT fibers to converge at the axial focal range of the Vevo PA system while not being blocked by the transducer itself, making simultaneous PDT and US-PAI possible as shown by the complete setup render in Fig. 1c. Therefore, adjusting the transducer to align the tumor within the PA focal range automatically aims the PDT fibers for optimal illumination.

### Synthesis of Liposomal Formulation of Benzoporphyrin Derivative

Liposomal benzoporphyrin derivative (BPD) was synthesized using a thin film hydration and extrusion method. As previously reported by us,^23,44,45^ the lipids cholesterol (ovine) (20 μmol), 1,2-dipalmitoyl-sn-glycero-3-phosphocholine (2.5 μmol), 1,2-dioleoyl-3-trimethylammonium-propane (chloride salt) (1 μmol), and 1,2-distearoyl-sn-glycero-3-phosphoethanolamine-N-[methoxy (polyethylene glycol)-2000] (ammonium salt) (10 μmol) (Avanti Polar Lipids) along with BPD (MedChem Express) (0.25 μmol) in chloroform (Thermo Fisher Scientific) were added to the Pyrex tube. The chloroform solution was evaporated under a gentle stream of nitrogen. Once the film was completely dry, it was hydrated with 1 mL of Dulbecco’s PBS (Thermo Fisher Scientific), vortexed for 5 seconds, then left in the hot water bath at 45°C for 10 minutes. The liposomal BPD solution was then vortexed vigorously and put on ice for 10 minutes. The freeze-thaw cycles were repeated for a total of 5 cycles. After the last cycle, the solution was extruded at 50°C through a 0.1 mm polycarbonate membrane (Cytiva Life Sciences) for a total of 6 cycles. Liposomes were then diluted 200x in PBS for dynamic light scattering and zeta potential measurements (Brookhaven Instruments). The average diameter of the liposomes obtained was 116.2 ± 13.2 nm with a polydispersity index of 0.24 ± 0.03 and a zeta potential of 20.9 ± 5.1 mV.

### Photodynamic Therapy (PDT)

The treatment groups had 100 mW/cm^2^ (n=5) and 400 mW/cm^2^ (n=5) laser irradiation, referred to as the low fluence rate (LFR) and high fluence rate (HFR) groups respectively. The control group (n=4) termed as “light only” received the equivalent higher fluence rate with no PS administered. Laser fluence rates were measured at comparable fiber-to-tumor distance and averaged across the four fibers for every case. Mice received liposomal BPD (0.5 mg/kg BPD eq.) intravenously via the tail vein and were kept in the dark until the irradiation time. After a 90-minute drug-light interval,^43^ the tumors were irradiated with a 690 nm continuous wave laser (Modulight Corporation) through a custom-designed four-way source-split optical fibers (Thorlabs, Inc.) mounted with our 3D printed fiber holder attachment. The tumors were treated for 15 minutes in each case, with simultaneous 2D US-PAI monitoring. A 5-minute baseline scan was conducted prior to irradiation and for 5-10 minutes following the cessation of the light interval.

We designed and 3D printed an additional PDT fiber holding setup with analogous fiber orientation geometry that allows a 12 × 12 mm square optical sensor (Ophir Starbright) to be rotated around a central normal axis, collecting individual fluence rates of each fiber at distances 22.5 mm (fiber holders along minor elliptical axis) or 25 mm (fiber holders along major elliptical axis). The laser current was adjusted until the total reading of the four PDT fibers was within 5 mW/cm^2^ of the target fluence rate for each respective treatment group.

### Ultrasound-Guided Photoacoustic Imaging and related Data Processing

US-PAI acquisition was conducted with the Vevo LAZR-X system (FUJIFILM, VisualSonics, Inc.) equipped with the MX250S 21 MHz LAT and a Nd:YAG nanosecond pulsed laser with a 20 Hz pulse repetition frequency. Hemodynamic images for StO_2_ were acquired with the Oxy-Hemo mode (750/850 nm PA acquisition) with maximum persistence (20 averages per frame) at a US gain of 22 dB and PA gain of 45 dB. Volumetric tumor scans were taken, at 0.152 mm step size to satisfy Nyquist criteria in the elevational direction, approximately 2 hours prior to and 30 minutes post-PDT. Changes in StO_2_ were monitored throughout PDT by acquiring PA images of the center cross-section of the tumor. With averaging, the resulting StO_2_ image temporal resolution was approximately 4 seconds.

B-scan US-PAI images during PDT, as well as 3D pre- and post-PDT images, were exported to accompanying Vevo Lab software (FUJIFILM, VisualSonics, Inc.), and tumor regions of interest (ROI) were annotated from the US images. The averaged StO_2_ and HbT values of the tumor region, excluding zero values, in every frame were exported to MATLAB (MathWorks, Inc.) as ‘.csv’ files, graphed as a function of time, and filtered with 15^th^ order ‘medfilt1’. StO_2_ values were normalized by applying a scale factor to the data calculated by dividing 100% by the mean StO_2_ value for the 5-minute pre-PDT 2D acquisition. 3D ROIs for pre- and post-PDT US-PAI images were rendered on the Vevo LAB software while average 3D tumor StO_2_ was exported for statistical analysis.

During the PDT process, either the PS or the oxygen is consumed. In our particular case, the point at which we see oxygen consumption not depleting any further is defined as the endpoint of “active PDT”. Normalized StO_2_ change rates for statistical analysis were computed by applying the ‘movingslope’ function to the filtered and normalized StO_2_ values over the first 5 minutes of PDT irradiation and averaging those values for each mouse, since the endpoint of active PDT varies between individuals. To generate frame LStO_2_ images, beamformed and spatially co-registered 2D US and PA images were exported to MATLAB and annotated with the Photoacoustic Annotation Toolkit for MATLAB.^46^ The tumor ROI was manually annotated from the US image. The 750 and 850 nm PA images were then spectrally unmixed using the MATLAB Optimization Toolbox function ‘lsqnonneg’ to obtain values of StO_2_ and HbT. The molar extinction coefficients of Hb and HbO_2_ used for spectral unmixing were taken from literature.^47^ A noise threshold was determined for each mouse by taking the mean HbT value of a 20 × 20-pixel area from both top corners of each frame in the 2D scan. Pixels where HbT was less than that value were set to zero for both the HbT and StO_2_ images. The LStO_2_ images were calculated from element-wise subtraction of the StO_2_ image at the end of ‘active PDT’ from the StO_2_ image immediately before light dose initiation. A 2D median filter using a 3x3 neighborhood was applied to each image for display purposes.

### Immunohistochemistry

Cryo-sections of the tumor (10 µm thick in the same orientation as PA images) were obtained utilizing a cryotome and fixed on glass microscope slides. Post-fixation, hematoxylin and eosin (H&E) staining was performed as previously described by Xavierselvan et al.^44^ The successive slides to the H&E slide were used to perform immunofluorescence (IF) staining. For IF staining, the cross sections were fixed using an acetone and methanol mixture (1:1). The sections were air-dried for 30 minutes post-fixation and then subjected to three sequential 5-minute washes with PBS. The sections were then blocked with 1% bovine serum albumin solution (Cat: 37525; Thermo Fisher Scientific) for 1 hour at room temperature. Primary antibody for targeting vasculature (Mouse CD31/PECAM-1 Polyclonal Goat IgG, Cat: AF3628; R&D Systems, 1:5 dilution) and antibody against pimonidazole adducts (conjugated IgG1 rat monoclonal antibody clone 11.23.22.r, Cat: Red 549 Mab; Hypoxyprobe, 1:25 dilution) were added to the sections and incubated overnight at 4 °C. Later, the primary antibody was washed off with PBS and secondary antibody (NorthernLights™ Anti-goat IgG-NL637 Cat: NL002; R&D Systems, 1:50 dilution) was added to tissue sections and incubated for 2 hours at room temperature. The sections were then washed in PBS and the nuclei were counterstained and mounted with 4’,6-diamidino-2-phenylindole (DAPI) SlowFade Gold Antifade Mountant (Thermo Scientific Cat: S36939).^48^ The stained slides were imaged with 20x objective using the EVOS M7000 imaging system (Thermo Fisher Scientific) with appropriate EVOS light-emitting diode (LED) excitation and emission filter cubes. IF and H&E images were processed in FIJI for display purposes. H&E images were white-balanced for display. The skin and background were cropped in the IF image and the rolling ball algorithm with a 500-pixel size radius was used to perform background subtraction. Additionally, brightness and contrast were adjusted to highlight high signal areas.

### Statistical Analysis

GraphPad Prism (La Jolla, CA, USA) was used to perform statistical analyses. Ordinary one-way ANOVA with Tukey’s multiple comparisons was used to compare the average normalized change in StO_2_ values in the tumor during the active PDT period across different PDT doses. A p-value < 0.05 is considered statistically significant.

## Results and Discussion

### Hemodynamic PA images obtained during PDT correlate with immunohistochemistry

Immunohistochemistry is considered a gold standard method to validate the oxygenation status of tumors where CD31 for blood vessels, pimonidazole or carbonic anhydrase stain for hypoxia are widely utilized.^49^ For example, we and others have previously demonstrated that tumor vasculature measured with PA imaging had good correlation with histological markers for vasculature (CD31).^37,50-52^ Gerling et al. and Tomaszewski et al. have shown that tumor StO_2_ measured with PA imaging has good correlation with the pimonidazole histological marker for regions lacking molecular oxygen.^3,31^ Pimonidazole tissue binding has been shown to rapidly increase at oxygen levels below 10 mmHg (∼10% StO_2_ per the Severinghaus Equation)^53^ and was used to spatially compare pockets of hypoxia shown in PA imaging. Similar to the previous studies, here we showcase the correlation between PA images and the immunofluorescence images from representative mice with FaDu tumors in Fig. 2 at varying degrees and spatial localizations of hypoxia.

**Figure 2:**
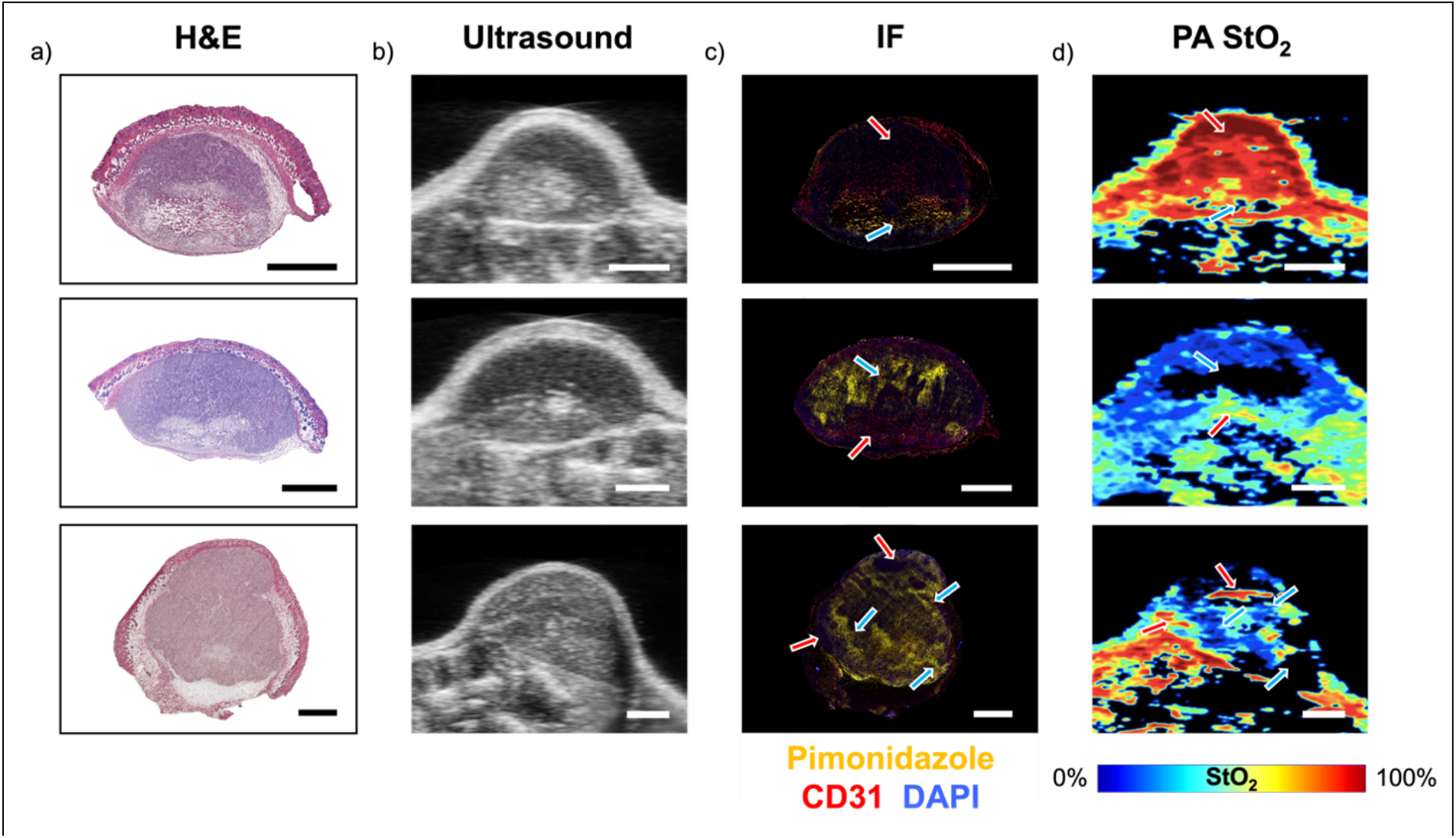
a) H&E and b) US B-scan images show structural similarities in the tumor while c) immunofluorescence images with pimonidazole stain (yellow regions) correlates to low-StO_2_ regions in d) PA images for 3 distinct tumors. Red arrows highlight areas of correlation between high StO_2_ and low pimonidazole signal while blue arrows highlight areas with low StO_2_ and high pimonidazole signal. Scale bars represent 2 mm.

The H&E images in Fig. 2a show structural similarities that match with the US images in Fig. 2b. The tumors are heterogeneous in morphology as well as tissue density, which provide varying contrast in both H&E and US. For example, the tumor in the top row of Figure 2, show cases two distinct areas in US image, hypoechoic top region and hyperechoic bottom region. These regions appear on the H&E image as regions with different cellular density, The IF images in Fig. 2c depict pimonidazole for hypoxia (yellow), CD31 for vasculature (red), and DAPI as a nuclear counterstain (blue) which are compared with the PA StO_2_ images in Fig. 2d. Highly oxygenated regions are represented by red in PA images while deoxygenated regions are represented by dark blue and black, with black indicating avascularity. Red arrows in Fig. 2c-d show matching non-hypoxic regions (low or no pimonidazole areas) while blue arrows highlight areas of hypoxia (high pimonidazole areas). The tumor in the top row shows overall high oxygenation in the PA image except for a small area at the base of the tumor. This is matched by an overall lack of pimonidazole stain in the tumor besides this small area at the base. In contrast, the tumor in the middle row shows major hypoxia in the upper majority of the tumor which corresponds to a large amount of pimonidazole in the top region of the tumor. The tumor in the bottom row has highly heterogeneous distribution of oxygen, likely related to its unique morphology that is once again matched by the pimonidazole stain. These regional variations of oxygenation may be caused by a combination of dysfunctional tumor vasculature, acute hypoxia following oxygen consumption during PDT, as well as hypoxia induced by PDT vascular damage. Besides validating our PA images, these results highlight the spatially varying oxygenation present in tumor tissue and therefore a significant need for spatially resolved oxygen imaging for PDT. One-dimensional sensors are simply insufficient to characterize the heterogenous oxygen status of a tumor before, during, or after treatment, and we thereby see significant motivation to bring US-PAI into the real-time image-guided PDT dosimetry field.

### US-PAI can Monitor Acute Oxygen Depletion in Real-Time During PDT

Figure 3 showcases the spatially and temporally resolved hemodynamic changes during PDT obtained with our custom built US-PAI transducer PDT fiber holder attachment. The graph in Fig. 3a depicts the averaged, normalized tumor StO_2_ over the duration of PDT and beyond. An ROI of the tumor was drawn from the US image (Fig. 3b, white ROI). From PA StO_2_ images, the average value of the pixels in the ROI were plotted against time. These values were also normalized by setting the average StO_2_ obtained prior to the PDT treatment to 100% and scaling the rest of the data accordingly. The normalization accounted for variation in the initial StO_2_ status of the tumor when comparing various mice and treatment regimens. The PA StO_2_ frames at critical time points during PDT are shown in Figs. 3c-h. while Video. S1 shows the images acquired during the entire 30 minutes.

**Figure 3:**
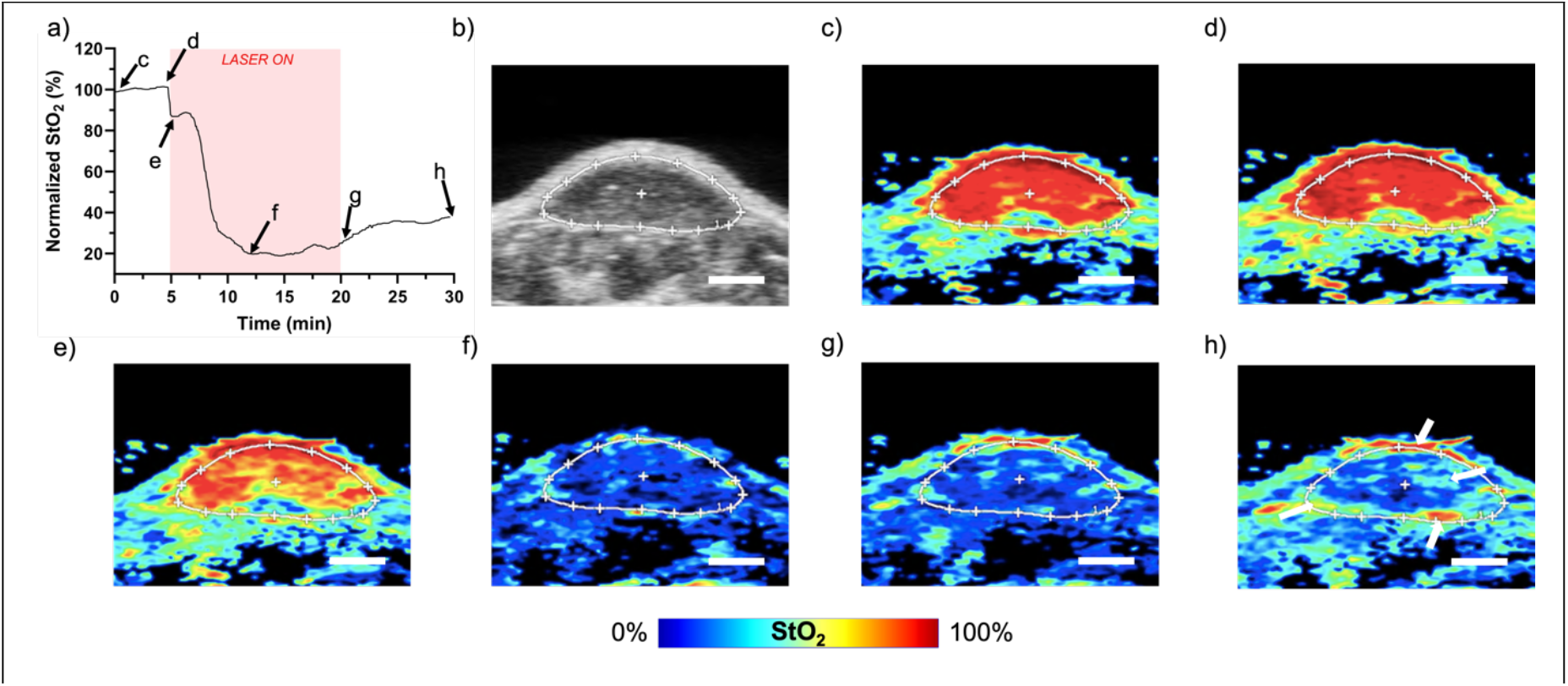
a) Normalized StO_2_ over the PDT timeline within the white ROI from the b) US B-scan image of the tumor. c-h) Corresponding PA StO_2_ images for time points annotated on the graph in (a). White arrows in (h) highlight areas of re-oxygenation. Scale bars = 2 mm. Video. S1 shows the movie of all PA StO_2_ images captured pre, during, and post-PDT. Red represents highly oxygenated areas in the PA StO_2_ images while blue represents deoxygenated regions.

Averaging the tumor region StO_2_ values, as shown in Fig. 3a, undermines the regional variation in tumor oxygenation that PA imaging is particularly adept at monitoring. Nevertheless, the average StO_2_ within the tumor region is a useful parameter in evaluating bulk PDT hemodynamic trends and comparing the results obtained previously with other 1D monitoring techniques. Within seconds of irradiating the tumor, a sharp decrease in StO_2_ is observed from Fig. 3d to Fig. 3e which we attribute to rapid ROS formation from accumulated PS consuming oxygen. After this initial drop, StO_2_ briefly rises which is consistent with previous studies. Vasodilation has been observed at the onset of PDT^54^, potentially due to thermal effects or a physiological response to the initial oxygen reduction.^55^ This results in an increase in blood flow at the beginning of PDT has been observed in many previous studies.^28,29,56,57^ The extra influx of oxygenated blood hinders, or in this case, reverses, the decline in StO_2_ from PDT oxygen consumption, which has been previously observed with diffuse reflectance spectroscopy^32^ and BOLD MRI monitoring.^33^ Following this increase in StO_2_, a steady decline in StO_2_ associated with the acute hypoxia generation from the primary PDT response is expected for an HFR individual.^58^ The decline of StO_2_ due to ROS formation is further enhanced by the decrease of blood flow resulting from acute vascular shutdown.^28^ Approximately 7 minutes into treatment at the PA frame shown in Fig. 3f, we observe that StO_2_ reaches a minimum and slowly begins to increase despite the continued irradiation of the tumor, indicating cessation of photodynamic action. We refer to this point where StO_2_ doesnot decrease further but is either stagnant or increase as the “end of active PDT” time point.

The hemodynamic response of the tumor to PDT is not uniform as seen in Figs 3c-h and Video. S1, emphasizing the need for spatially resolved real-time monitoring and clearly demonstrating the downside of 1D bulk tissue measurements. The re-oxygenation shown in the Fig. 3a graph from 12 to 30 minutes is not representative of the entire tumor image as this effect is primarily isolated to the tumor rim as well as an area in the top-right quadrant of the cross section. The white arrows in Fig. 3h point to these regions. While the reason for the heterogeneous response could be insufficient dose due to a number of factors such as regionally varying vascular status, PS localization, or tissue optical properties, the ability to temporally and spatially monitor oxygen-utilizing cancer therapies is an important aspect of dose personalization and we demonstrate in the following sections that PA imaging is able to decipher between lower and high fluence rates.

### US-PAI shows Fluence Rate Dependent Oxygen Depletion Rates

In Fig. 4, we showcase the data that real-time US-PAI monitoring can capture differences in oxygen utilization at different dosing regimens. The graph in Fig. 4a again shows the bulk normalized StO_2_ in the tumor region before, during, and after PDT average for several mice while Fig. 4b showcases data from one representative mouse in the LFR (blue line), HFR (red line), and light only t (green line) treatment groups. The first noticeable difference between the HFR and LFR individuals is the high StO_2_ decrease in HFR group compared to the LFR group. This decrease also happened at a much faster rate in the HFR group. The overall rate of decline in the HFR individual is greater than the LFR individual, and the overall decrease of StO_2_ is greater for the HFR individual due to the well-established oxygen conservation in LFR PDT.^19,59^ Blood vessels still receive the physiological perturbation from the onset of irradiation at LFR, but with PDT happening at a decreased fluence rate, the photodynamic action occurs at a slower rate where the onset of acute vascular shutdown is delayed and the vessels stay dilated for longer.^19,29,60,61^ The LFR group showed a statistically insignificant depletion rate of StO_2_ in the first 5 minutes of irradiation compared to the control while the HFR group shows a very high depletion rate, highly significant from both the LFR and light-only sham groups.

**Figure 4.**
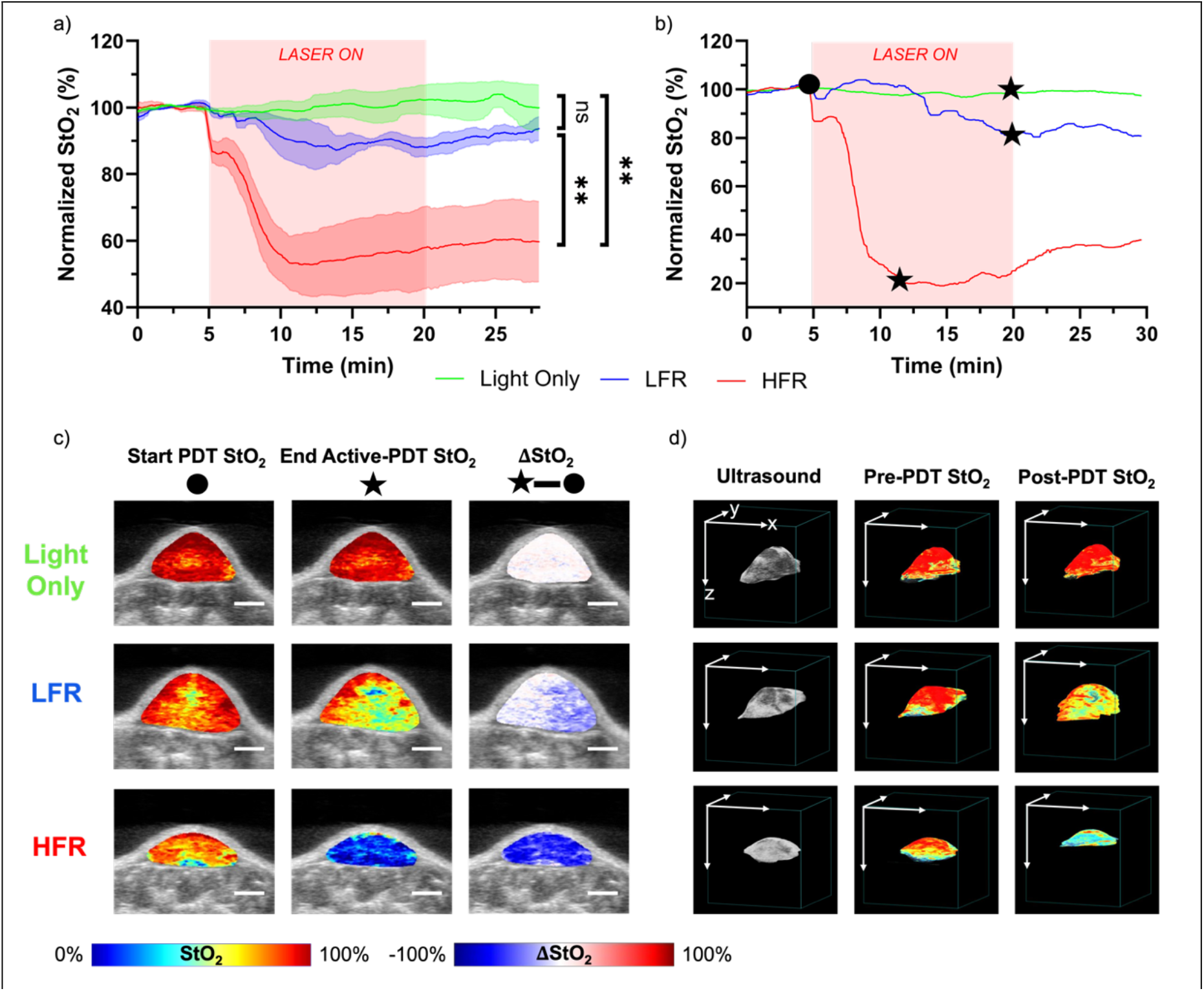
a) Normalized StO_2_ over the PDT timeline averaged with SEM for all mice in LFR, HFR, and light only treatment groups. Significance difference was measured by comparing the average rate of StO_2_ change during the first 5 minutes of light irradiation using one-way ANOVA with Tukey’s post-hoc test (ns-not significant; **p-value < 0.01). b) Representative traces for one tumor in each light dose group from (a). The black dot represents the start of PDT while the black star represents the end of active PDT for each mouse. c) 2D US-PAI immediately before and after active PDT, as well as their difference maps. Scale bars = 2 mm d) 3D US, as well as pre- and post-PDT PA StO_2_ tumor renders. Axis scale bars = 10 mm. For StO_2_ 2D and 3D images, the red in the colorbar represents highly oxygenated areas while blue represents deoxygenated regions. For ΔStO_2_ images, the red in the colorbar represents increased oxygenation regions, blue represents decreased oxygenation regions, and white indicates no change.

Focusing on 3 representative treatment group individuals in Fig 4b, it is shown that active PDT lasts until the end of the light dose for the LFR individual, unlike the mouse in the HFR group. The end of active PDT is denoted on the graph with a black star for each StO_2_ trace, and the StO_2_ PA images in the left and middle column of Fig. 4c show the frames at the beginning of the light dose (black dot) and the end of active PDT (black star) for each individual, respectively. The right column of Fig. 4c shows the difference in these images, signifying the spatially-resolved extent of deoxygenation over the active PDT period. It is intuitive that a faster delivery of the light dose will lead to a greater overall deoxygenation over a shorter timeframe in the HFR individual over the LFR individual, but it is again apparent that these changes do not happen uniformly. In addition, it is well-accepted that HFR PDT is self-inhibitory as oxygen is removed by ROS faster than it can be resupplied by the bloodstream.^61^

The 3D renders in Fig. 4d reflect the US, pre- and post-PDT PA images of representative mice in different treatment groups. Specifically for the HFR tumor, a gradual reoxygenation post-PDT was observed. But the 3D scans post-PDT donot capturethe maximum hypoxia that was present during and nearer to the end of treatment. For most of the tumors, light dose-dependent decrease in StO_2_ was observable with maximum decrease observed in the HFR group. It is to be noted that, across the study, the extent of re-oxygenation after active-PDT varied amongst different tumors in the LFR and HFR groups. However when the heterogenous StO_2_ values within a specific tumor were averaged for the entire tumor and plotted as a single data point, no statistically significant difference was observed between pre-PDT and immediately post-PDT StO_2_ values, agreeing with the previous findings of Mallidi et al.^23^ With the tumor microenvironment changing so rapidly in the immediate minutes following PDT, heterogenous hemodynamic information is lost by not conducting real-time monitoring.

### Post-Active PDT Reoxygenation is in agreement with Blood Vessel Perfusion Immunohistochemistry

It has been established previously by several studies through histology^54^, PA microscopy^62,63^ or intra-vital microscopy^64^ that vessel damage is heterogenous within the tumor. In Fig. 5 we demonstrate that change in PA StO_2_ maps post active PDT are in good agreement with these previous observations that vascular damage is heterogenous within the tumor. Specifically, immunofluorescence images of vascular perfusion observed through overlay of TL (vascular perfusion) and CD31(tumor vasculature) was compared to change is PA StO_2_ maps (reoxygenation) calculated from the images obtained at the end of active PDT to the end of the monitoring period. The change in StO_2_ image shown in Figs. 5a and 5d was overlaid on US gray scale and pseudocolored where positive change represented by red indicated significant reoxygenation in the tumor while blue represented the opposite effect. Figs. 5b, 5c, 5e and 5f showcase IF stains of two representative tumors, where TL labeling perfusive vasculature is shown in green, and CD31 labeling all vasculature is depicted in red pseudocolor. Overlap of TL with CD31 indicated the presence of functional vasculature. The two insets in Figs. 5a and 5d represent an an area of re-oxygenation (ROI with red box) and an area of minimal re-oxygenation (ROI with blue box). Corresponding vascular IF images for the insets are found in Fig. 5b-c for the tumor in Fig. 5a, and Fig. 5e-f for the insets in Fig. 5d. A prevalent CD31 stain, indicative of highly vascularized tumors, can be observed in all the IF images of Fig. 5. On the other hand, TL signal is prevalent only in Fig. 5b and 5e that corresponds to regions of re-oxygenation post-active PDT. Vascular regions that had increase in oxygenation must retain some level of functionality following treatment, potentially indicating areas of sub-optimal dosing and incomplete or no vessel damage. Identifying areas of re-oxygenation in the entire tumor depth following the end of active-PDT can add significant dosimetry value. In prior studies, changes to vascular function has been monitored with limited temporal or spatial resolution during PDT using PA microscopy or intravital imaging that are capable of resolving individual blood vessels, but lack sufficient penetration depth.^63^ With availability of mesoscale US-PAI demonstrated here, we are able to obtain, pre-PDT vascular status of the tumor, post-PDT status immediately post irradiation or several hours post irradiation to decipher the heterogenous PDT-induced dysfunctional vasculature. Our future studies will involve quantifying and evaluating the heterogenous areas of re-oxygenation as surrogate marker for long-term tumor response to treatment.

**Figure 5:**
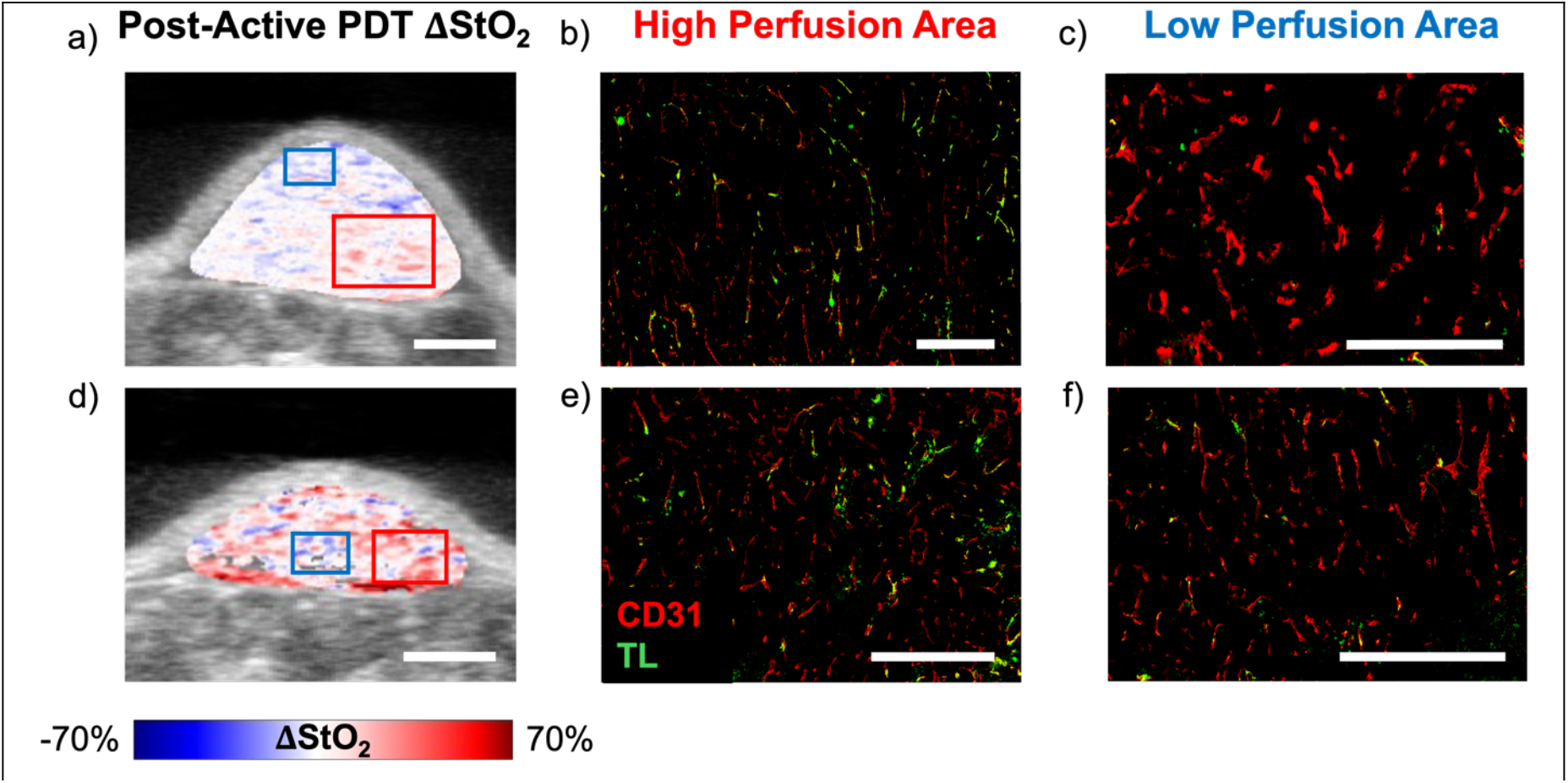
a) Map of the change in PA StO_2_ from the end of active PDT to the end of the monitoring period overlaid on US. Red corresponds to areas of re-oxygenation while blue corresponds to areas of deoxygenation. Scale bars = 2 mm. IF images with CD31 (vasculature) and TL (perfusive vasculature) b) correspond to the red inset of re-oxygenation in (a) where TL-stained vessels are prevalent and c) regions of deoxygenation/no change where vessels are mainly unstained by TL. Scale bars = 500 µm. d-f) Repeat of (a-c) for another mouse tumor.

### Spatial Resolution of US-PAI can Detect Heterogenous Dosing

A major advantage of US-PAI over other previously used methods is its spatial resolution and the resulting ability to detect heterogeneity in treatment response. Thus far we have successfully demonstrated that US-PAI is a feasible method to monitor StO_2_ in real-time during PDT and the effective administered PDT dose is inherently heterogeneous due to the combination of factors previously discussed. Here we aimed to exaggerate this dosimetry disparity by modifying the configuration of the light dose. For the HFR individual featured in Fig. 6, the two PDT optical fibers directed towards the face of the imaging plane were kept at 100 mW/cm^2^ while the fiber to the left of the imaging plane was increased to 150 mW/cm^2^ and the fiber to the right was decreased to 50 mW/cm^2^ as shown by the schematic in Fig. 6a. The averaged StO_2_ graph in Fig. 6b also contains traces for the averaged and normalized bulk StO_2_ values in the left and right sides of the tumor, while the US image in Fig. 6c shows the overlayed left (magenta ROI) and right (cyan ROI) half of the tumor. The H&E stain shown in Fig. 6d has similar hemispherical the structural features as in the US image. Selected PA StO_2_ images at the same cross-section as the US image are shown in. All the images obtained pre and during PDT regimen are shown in Video. S2. Clearly images in Fig. 6e-j show the tumor deoxygenate substantially in the left half of the tumor that received higher fluence rate and light dose. The tumor consequently re-oxygenates more on the left side after the end of active-PDT at the frame shown in Fig. 6m. Interestingly, the corresponding TL stain in Fig. 6l shows that most blood vessels remaining functional at the end of treatment is largely localized to the left side of the cross-section where there was a greater light dose and re-oxygenation. In other words, despite having a higher light dose, more vasculature remains functional on the left side of the tumor when compared to the right. While the light dose was increased, the effective PDT dose was diminished by hypoxia during much of the irradiation time. These findings are supported by the post-PDT US-PAI image in Fig. 6o, occurring 20-30 minutes following the end of the monitoring session and prior to euthanasia, which corresponds to the pimonidazole immunofluorescence image in Fig. 6n. The same cross-section shows hypoxia development mainly in the right side of the tumor, and more specifically, in regions that did not show prominent reoxygenation. This provides promise that tumor regions that don’t show re-oxygenation may be responding to PDT better than those that do re-oxygenate, given the connection of hypoxia development to PDT efficacy, despite the early time point.^23^ Furthermore these observations are in alignment with previous studies that demonstrated that higher fluence rates donot always lead to effective outcomes in PDT and that the vascular damage is higher with lower fluence rates.^19,54,64^ To our knowledge, an asymmetric light dose model for *in vivo* PDT experiments has not previously been investigated, but may provide utility in inducing controlled dose variation to test spatially resolved monitoring techniques for PDT monitoring such as US-PAI.

**Figure 6:**
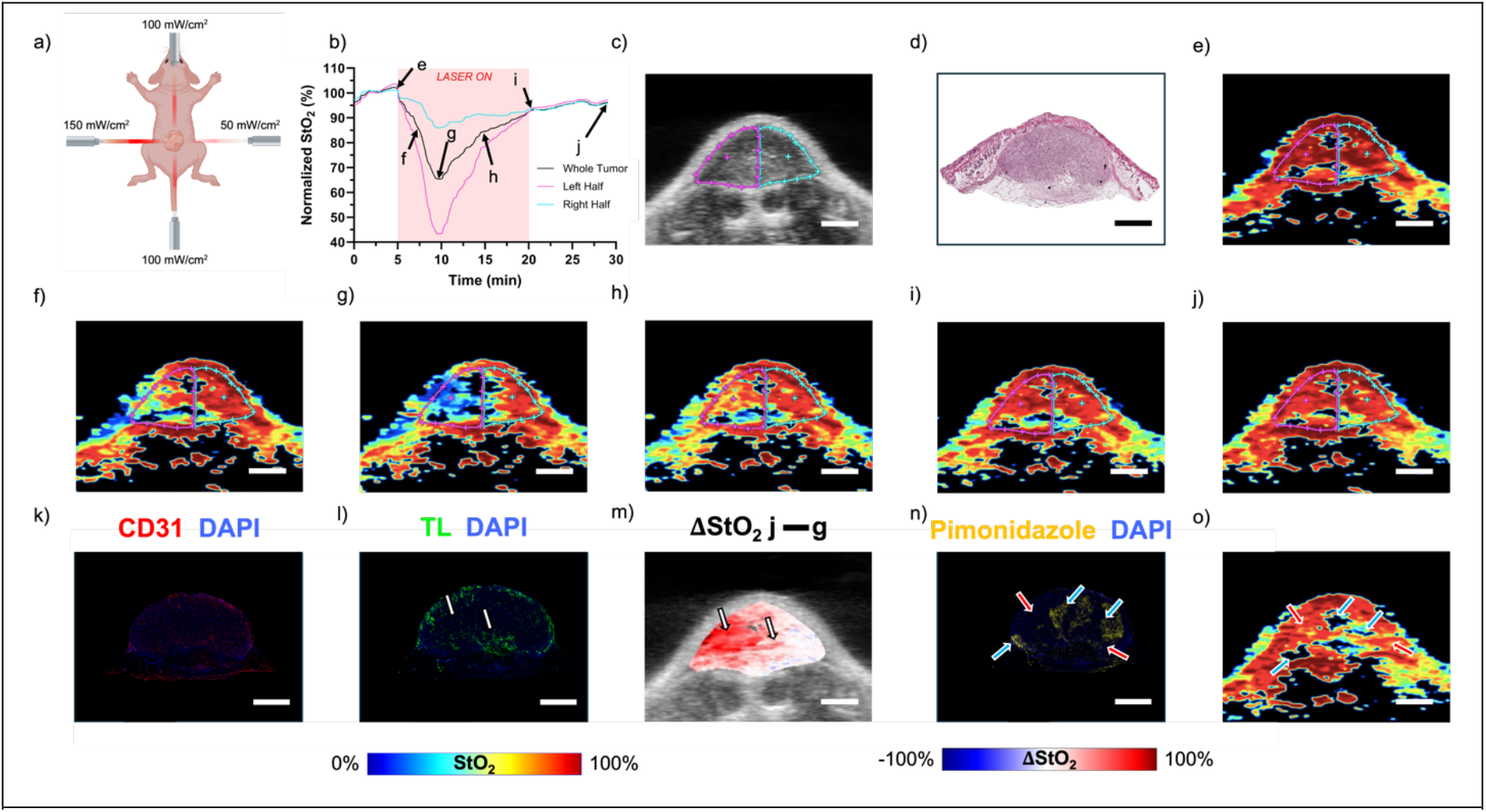
a) Schematic of the non-uniform light dose arrangement. b) Normalized StO_2_ over the PDT timeline in the ROIs for left half, right half, and whole tumor segments. c) US B-scan image of the tumor with left and right half ROIs and d) matching H&E histology image. e-j) Associated PAI StO_2_ images for annotated timepoints in b). k) CD31 and l) TL stains of the cross-section matching the PA image. m) Difference of frames g (end of active-PDT) and j (end of scan) showing areas of re-oxygenation. White arrows highlight regions of correlation between functional vessels in the TL stain and re-oxygenation in m. n) Pimonidazole matches with low-StO_2_ regions in o) the corresponding frame from the post-PDT 3D PA image. Red arrows highlight areas of correlation between high StO_2_ and low pimonidazole while blue arrows highlight areas with low StO_2_ and high pimonidazole. Scale bars = 2 mm. Video. S2 shows the movie of all PA StO_2_ images captured pre, during, and post-PDT.

## Conclusions and Future Directions

In this study, we have demonstrated the feasibility and utility of US-PAI for real-time StO_2_ monitoring of photodynamic action. Our custom-built probe integrating the PDT illumination fibers with the US-PAI transducer has enabled monitoring of hemodynamic markers throughout treatment, particularly spatially resolved heterogeneity in tumor oxygen utilization. Mapping of tumor re-oxygenation post-treatment to vascular perfusion, via IF staining, gave us an indication for how real-time PA StO_2_ data may aid in predicting responsive and non-responsive regions, and thereby treatment outcome. Photodynamic action is a complex phenomenon where the effective PDT dose is influenced by the spatial and temporal overlap of the PS concentration, PS molar extinction coefficient at the treatment wavelength, PS ROS quantum yield, light dose photon density, and the molecular oxygen concentration.^16^ While PA imaging has been previously used to map heterogenous PS uptake,^65^ recently pump-probe-based photoacoustic tomography has been empoloyed to evaluate partial pressure of oxygen within the tumors to aid in PDT dosimetry.^66,67^ However, this imaging technique requires sophisticated synchronization between expensive laser equipment as well as a very photostable dye such as methylene blue. Infact, methylene blue is the only dye that has shown success in pump-probe based photoacoustic tomography^66,67^ due to its strong absorption and low photobleaching, but lacks the potency of a high ROS generating PS like BPD. Our future studies will involve mapping of the heterogenous PS accumulation alongside StO_2_ maps to more holistically model the effective PDT dose required for personalized treatments. Additionally, further developement of the imaging equipment is required for clinical translation of the methodology presented here. It is to be noted that with the commercial US-PAI system used in this study, we were limited to 30 minutes of monitoring data due to the system’s buffer capacity. However, with the availability of customizable US-PAI devices^68-70^ where data storage can be independent of data acquisition, there is a possibility to enable monitoring of long PDT sessions such as those required for low irradiance or fractionated light doses, as well as extended post treatment monitoring. The availability of such custom built systems will also enable synchronization with PDT lasers, enabling real-time modulation of the light dose for fractionated PDT. Infact fractionated PDT regimen has more effective outcomes than continuous PDT irradiation as it allows the tumor to reoxygenate or facilitate more tumoral PS accumulation.^25,26,29^ Also, the sensing electronics for US-PAI can be compacted enough to integrate into an all-inclusive probe for large animal or human PDT applications. Advancements in highly portable, inexpensive LED-based US-PAI systems^37,71^ make this PDT monitoring method feasible for use in low resource settings. Our future PDT studies are geared towards incorporating PA imaging of PS along with StO_2_ changes in real time with low cost mobile LED based PDT and US-PAI devices. Finally, PDT is one of the many oxygen-utilizing cancer therapies that could potentially benefit from real-time monitoring, so an analogous design to ours could feasible be built and tested on other treatment methods such as radiotherapy.

## Abbreviations

BOLD fMRI: blood oxygenation level-dependent functional magnetic resonance imaging
BPD: benzoporphyrin derivative
DAPI: 4’,6-diamidino-2-phenylindole
DCS: diffuse correlation spectroscopy
FBS: fetal bovine serum
H&E: hematoxylin and eosin
HbD: deoxyhemoglobin
HbO: oxyhemoglobin
HFR: high fluence rate
HNC: head and neck cancer
IACUC: Institutional Animal Use and Care Committee
IF: immunofluorescence
LAT: linear-array transducer
LED: light-emitting diode
LFR: low fluence rate
PA: photoacoustic
PBS: phosphate-buffered saline
PDT: photodynamic therapy
PS: photosensitizer
ROI: region of interest
ROS: reactive oxygen species
StO_2_–: blood oxygen saturation
TL: tomato lectin
US: ultrasound
US-PAI: ultrasound-guided photoacoustic imaging

## Acknowledgements

The authors would like to acknowledge funds for Mallidi from NIH S100D026844, R01CA266701, R21CA263694, and Tufts University School of Engineering. Additionally, we would like to thank Dr. Christopher D. Nguyen for the render of the experimental setup as well as Aayush Arora, Deeksha Sankepalle, Brian Li, and Daniel Wong for assistance with the experiments.

## Declaration of Competing Interest

The authors declare no commercial or financial relationships that could be construed as a potential conflict of interest in the conduct of this research.

## Supplementary

|  |  |
| --- | --- |
| Video 1 | Movie of all PA StO<sub>2</sub> images captured pre, during, and post-PDT of the mouse tumor featured in Fig. 3, along with associated animated graph from Fig. 3a. |
| Video 2 | Movie of all PA StO<sub>2</sub> images captured pre, during, and post-PDT of the mouse tumor featured in Fig. 6, along with associated animated graph from Fig. 6b. |

## Notes

### Competing Interest Statement

The authors have declared no competing interest.

